# Contemporary circulating enterovirus D68 strains show differential viral entry and replication in human neuronal cells

**DOI:** 10.1101/331116

**Authors:** David M Brown, Alison M Hixon, Lauren M Oldfield, Yun Zhang, Mark Novotny, Wei Wang, Suman R. Das, Reed S Shabman, Kenneth L Tyler, Richard H Scheuermann

## Abstract

Historically, enterovirus D68 (EV-D68) has primarily been associated with respiratory illnesses. However, in the summers of 2014 and 2016 EV-D68 outbreaks coincided with a spike in polio-like acute flaccid myelitis/paralysis (AFM/AFP) cases. This raised concerns that the EV-D68 virus could be the causative agent of AFM during these recent outbreaks. To assess the neurotropic capacity of EV-D68, we explored the use of the neuroblastoma-derived neuronal cell line, SH-SY5Y, as a tissue culture model to determine if differential infection permissibility is observed for different EV-D68 strains. In contrast to HeLa and A549 cells, which support viral infection of all EV-D68 strains tested, SH-SY5Y cells only supported infection by a subset of contemporary EV-D68 strains, including members from the 2014 outbreak. Viral replication and infectivity in SH-SY5Y was assessed using four different assays – infectious virus production, cytopathic effects, cellular ATP release, and VP1 capsid protein production – with similar results. Similar differential neurotropism was also observed in differentiated SH-SY5Y cells, primary human neuron cultures, and a mouse paralysis model. Using the SH-SY5Y cell culture model, we determined that barriers to viral entry was at least partly responsible for the differential infectivity phenotype, since transfection of genomic RNA into SH-SY5Y generated virions for all EV-D68 isolates, but only a single round of replication was observed from strains which could not directly infect SH-SY5Y. In addition to supporting virus replication and other functional studies, this cell culture model may help confirm epidemiological associations between EV-D68 strains and AFM and allow for the rapid identification of emerging neurotropic strains.

**Author Summary:** Since the outbreak during the summer of 2014, EV-D68 has been linked to a type of limb paralysis referred to as acute flaccid myelitis (AFM), with evidence mounting for the causal link of EV-D68 to AFM. Among these AFM cases, concurrent EV-D68 infection was confirmed in several independent epidemiological clusters in four continents. In this report, we describe a neuronal cell culture model (SH-SY5Y cells) where only a subset of contemporary 2014 outbreak strains of EV-D68 show infectivity in neuronal cells, or neurotropism, based on four different assays of viral replication and infection. We further confirmed the observed difference in neurotropism *in vitro* using primary human neuron cell cultures and *in vivo* with a mouse paralysis model. Using the SH-SY5Y cell model, we determined that a barrier to viral entry is at least partly responsible for neurotropism. SH-SY5Y cells may be useful in determining if specific EV-D68 genetic determinants are associated with neuropathogenesis, and replication in this cell line could be used as rapid screening tool for identifying neurotropic EV-D68 strains. This may assist with better understanding of pathogenesis and epidemiology, and with the development of potential therapies.

## Introduction

The *Enterovirus* genus in the *Picornaviridae* family comprises many important human pathogens, including human rhinoviruses (HRV), the most common viral agents of the common cold; polioviruses, the causative agent of poliomyelitis; enterovirus A71 (EV-A71), associated with a variety of neurological diseases; and enterovirus D68 (EV-D68). Enteroviruses appear to continually circulate in human populations, with most infections being asymptomatic. For example, up to 72 percent of poliovirus infections are asymptomatic [1]. When poliovirus infections are symptomatic, they can cause a wide spectrum of clinically-distinct syndromes, ranging from minor, non-specific illness, to non-paralytic aseptic meningitis and flaccid paralysis [2]. Before the widespread use of effective vaccines, poliovirus-induced paralysis reached a peak of 21,000 cases in the U.S. in 1952 [3].

EV-D68 was first detected in children with pneumonia and bronchiolitis in 1962 [4]. Until recently, EV-D68 was one of the most rarely reported enteroviruses, with only 26 cases documented by the National Enterovirus Surveillance System in the U.S. from 1970 to 2005 [5]. Beginning in 2009, multiple contemporary clades began emerging worldwide [6]. In the summer and fall of 2014, 49 U.S. states experienced a nationwide outbreak of severe respiratory illness associated with EV-D68, with 1,153 confirmed cases, including 14 deaths [7]. Shortly after the U.S. outbreak, EV-D68 infections were also reported in Canada, Europe, and Asia. The total number of reported EV-D68 cases in 2014 exceeded 2,000 from 20 countries, resulting in the public health community classifying EV-D68 as a re-emerging pathogen of public health concern [8].

Reports of acute flaccid myelitis (AFM) occurring coincident to the outbreak of respiratory disease attributed to EV-D68 raised the possibility that EV-D68 might be the causative agent [7]. EV-D68 infection within a subset of these AFM cases was confirmed in several independent epidemiological clusters in the U.S. [9-14], France [15], Norway [16] Canada [17] and Australia [18]. Statistical analyses of the AFM cases in Colorado [12] and California [19] have supported the association between EV-D68 and AFM, and viral nucleic acid detection studies of patient samples have failed to reveal an alternative etiology [7, 10]. During the 2014 EV-D68 outbreak, patients presenting with AFM showed distinctive magnetic resonance imaging (MRI) findings characterized by brain stem and gray matter longitudinally extensive spinal cord lesions. This matches the findings described in previous outbreaks of EV-A71-associated AFM [9, 19, 20], suggesting that an enterovirus may be responsible. In support of this hypothesis, Hixon *et al.* [21] established that several contemporary EV-D68 strains, but not the historically archetypal Fermon and Rhyne EV-D68 strains, can cause a paralytic disease in neonatal mice due to viral infection and killing of spinal cord motor neurons.

Phylogenetic analysis reported that many of the 2014 EV-D68 outbreak isolates associated with AFM appeared to belong to the phylogenetic subclade, B1 [10, 22]. Interestingly, 12 substitutions identified in B1 2014 isolates carry the same amino acid or nucleotide residues observed at equivalent positions in other paralysis-causing enteroviruses, including poliovirus and EV-A71 [22]. This suggests that one or more of the nucleotide substitutions present in contemporary EV-D68 strains and lineages and not found in historical archetypal strains, may be responsible for the apparent increased incidence of neuropathology associated with the 2014 outbreak. EV-D68 has continued to evolve since the 2014 outbreak, which is unsurprising as mutation and recombination are known to occur in enteroviruses [23, 24]. Sequence analysis has led to the classification of a new clade D (a subclade of A) [25, 26], and a new subclade, B3, has emerged and quickly expanded [25, 27, 28]. Neurological symptoms have been associated with the novel B3 clade in Sweden [29], the Netherlands [30], Taiwan [31], Italy [32] and the United States [33] which experienced another AFM outbreak during the 2016 enterovirus season (summer and fall), with a total of 149 confirmed cases [2]. The seasonality and magnitude of this AFM outbreak matches the AFM surge observed in 2014. Additional surveillance of potentially emerging neurotropic or neuropathogenic strains is warranted.

To test if a specific genotype is associated with neurological symptoms, we report the development of a cell culture infection model based on the neuroblastoma cell line SH-SY5Y, that shows differential infectivity by different EV-D68 isolates. We observe a correlation between infection and replication in SH-SY5Y cells and neuropathogenesis in mice. This neuronal SH-SY5Y model may be useful for analysis of virus-host interactions *in vitro* and provides a facile assay to quantify which EV-D68 strains are neurotropic and neuropathogenic, potentially leading to better surveillance of virulent EV-D68 strains.

## Results

### SH-SY5Y cells express higher levels of neuron-specific genes than other candidate cell lines

Seeking a human cell culture to model neuron-specific infectivity, we performed expression profiling of two commonly used ‘neuronal-like’ cell lines, SH-SY5Y and HTB10. These cell lines were compared with HeLa cells as a non-neuronal permissive cell culture model. Both of these neuronal cell lines were first cultured by Biedler *et al.* in the early 1970s. SH-SY5Y is a subclone of the HTB11 (SK-N-SH) neuroblastoma cell line that was selected as an apparently homogenous population of cells with neuronal cell morphology. HTB10 (also known as SK-N-MC), reported as a neuroepithelioma cell line, has been used as a model for different neurotrophic viruses, such as hepatitis C poliovirus [34, 35] and enterovirus A71 [36]. We used RNA sequencing (RNAseq) to determine the genes expressed in SH-SY5Y, HeLa and HTB10, and specifically assessed the expression of twenty-six neuronal cell marker genes that were selected from the Allen Brain Atlas [37-39] (http://brain-map.org) and from BioGPS [40, 41] (http://biogps.org) as being highly neuron-specific (**Table 1).** Of the 26 selected genes, 22 showed measurable expression in SH-SY5Y cells, 21 of which showed higher expression levels in SH-SY5Y compared to HTB10 cells, and little if any expression in HeLa cells. These findings support the use of SH-SY5Y as a model neuronal cell line, while raising questions about the suitability of HTB10 as a “neuronal-like” cell line.

**Table 1.**
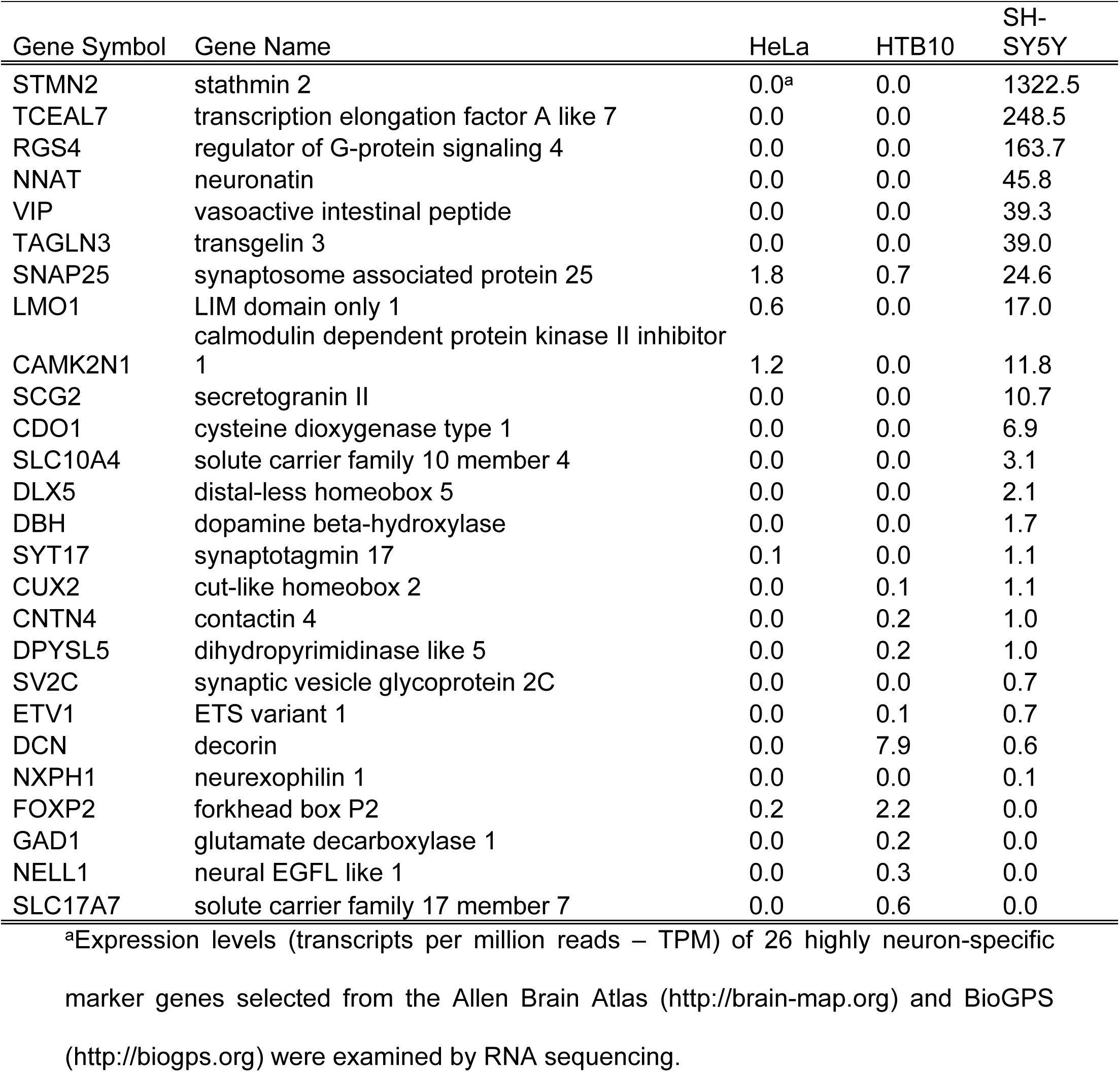
Expression of neuron-specific genes in HeLa, HTB10 and SH-SY5Y cell lines

### A representative B1 clade EV-D68 strain, US/MO/47, replicates in SH-SY5Y cells

Given the epidemiological association of recent EV-D68 infections with AFM, we sought to determine if there are any differential growth phenotypes between contemporary and historical EV-D68 strains in neuronal versus non-neuronal cell lines. To measure viral replication kinetics, each cell line was infected at a multiplicity of infection (MOI) of 0.1 and the virus growth was measured by determining virus titers (TCID_50_) in culture supernatant at five time-points post-infection. To examine whether viruses from the B1 clade showed any phenotypic differences in their ability to infect human neuronal cells, we first selected three different viruses to represent the phylogenetic diversity of EV-D68. US/MO/14-18947 from Missouri (US/MO/47) was selected as a representative of the B1 clade since it carries all 21 substitutions identified in our previous comparative genomics analysis [22]. USA/N0051U5/2012 from Tennessee (US/TN) was selected as a representative of clade A since it was isolated in the U.S. during roughly the same timeframe as US/MO/47 and possesses none of the 21 substitutions. VR1197 was selected as an example of an historical isolate similar to the prototypical Fermon strain isolated in 1962.

All three EV-D68 strains replicate in the non-neuronal cell lines HeLa (**Fig 1A**) and the alveolar A549 cell line (**S1 Fig**). These viruses also cause cell death as judged by visual evidence of cytopathic effects (CPE) in infected cell culture (**Fig 1B**). In contrast, only the US/MO/47 could replicate in the SH-SY5Y neuronal cell line, reaching peak titers of ∼10^5^ TCID_50_/ml by 48 hours post-infection (hpi). US/TN and VR1197 did not show any signs of replication, with titers not exceeding background after 96 hpi (**Fig 1A**). Similarly, CPE was observed after infection of SH-SY5Y cells with US/MO/47, but not with US/TN or VR1197 (**Fig 1B**). Similar results were seen at MOIs of 0.01 and 1.0 (**S2 Fig**). Furthermore, multiple passages of US/TN or VR1197 infected-supernatant onto fresh SH-SY5Y cells (passaging every 4 days for 12 days) failed to produce CPE. No increase in viral titers above background levels was detected by any virus following infection of HTB10.

**Fig 1.**
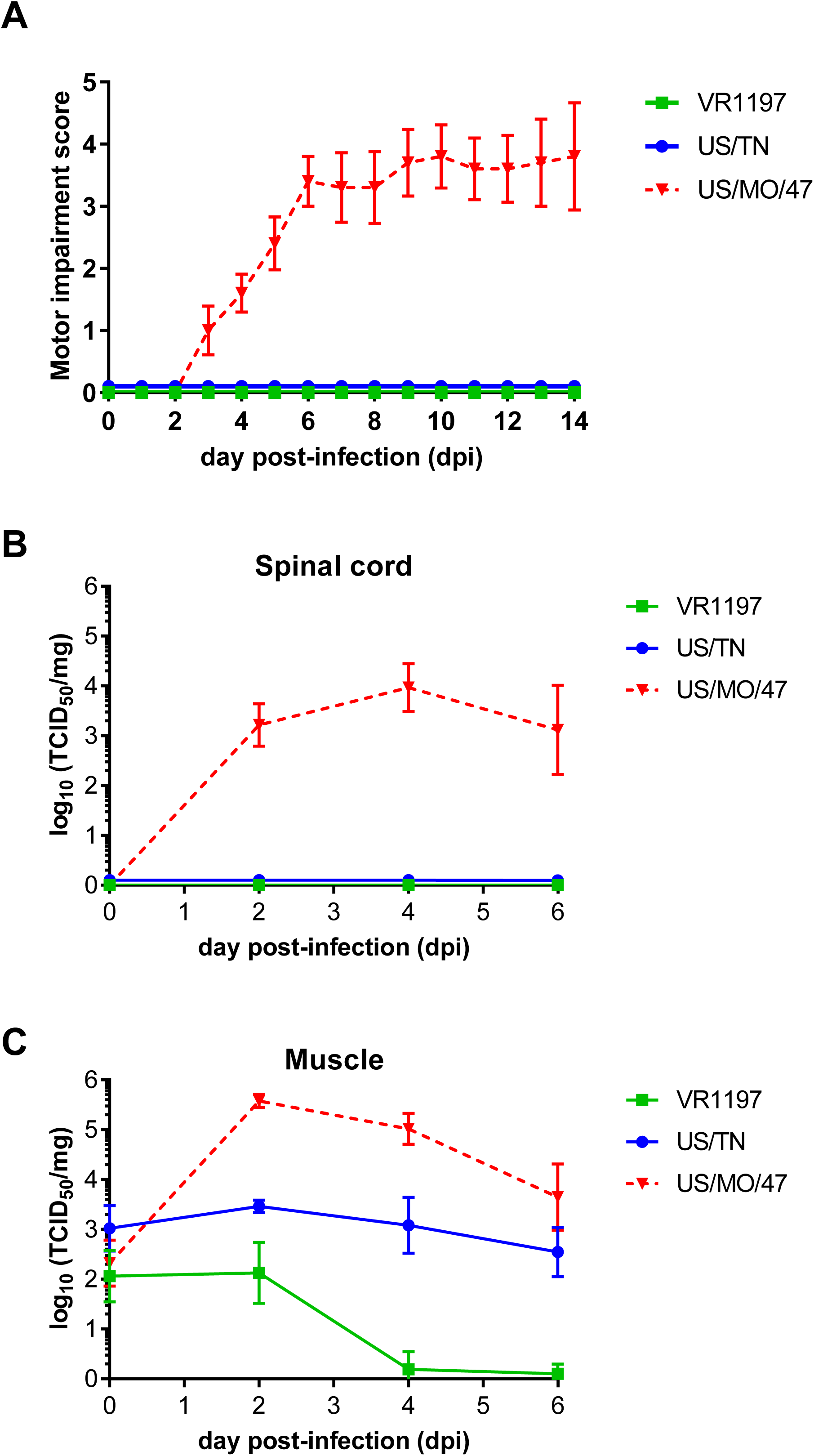
Differential infection and replication of EV-D68 strains in SH-SY5Y. A) SH-SY5Y, HTB10, and HeLa were grown to 90% confluence in 96-well plates before infection with EV-D68 US/MO/47, US/TN and VR1197 at a MOI = 0.1. Infection media was removed 2 hours post-infection (hpi) to reduce background. Cell culture lysates were collected at various time points after infection, and viral titers measured using endpoint dilutions for growth in HeLa cells. Dotted black line indicates the limit of detection. Error bars represent SEM of three biological replicates. B) SH-SY5Y, HTB10, and HeLa were infected with EV-D68 US/MO/47, US/TN and VR1197 at an MOI = 0.1 as above. Cells were visualized at 72 hpi with bright field microscopy at 400x. C) HeLa and SH-SY5Y cell lines were infected with the indicated EV-D68 strains at a MOI of 1.0. Cells were fixed at 18 hpi and stained with polyclonal antiserum against EV-D68 VP1 (red) and counterstained with DAPI (blue) for nuclei detection.

### Immunofluorescence confirms US/MO/47 replication in SH-SY5Y cells

We also examined production of the virus VP1 protein during infection of HeLa and SH-SY5Y cells. VP1 is among the initial proteins expressed following picornavirus infection, preceding capsid assembly [42]. We examined the ability of US/MO/47, US/TN and VR1197 viruses to synthesize the VP1 capsid protein 18 hpi using immunofluorescence. VP1 protein was detected in cells following infection of HeLa cells with all three EV-D68 strains (**Fig 1C**). However, only the US/MO/47 isolate produced VP1 following infection of the SH-SY5Y neuronal cell line (**Fig 1C**), which is consistent with our data on viral replication and CPE (**Fig 1A and 1B**). Also consistent with the viral replication experiments, no VP1 was produced by any of the three strains when HTB10 cells were infected. As typically observed in picornaviruses, VP1 staining was observed in the cytoplasm and not the nucleus of both HeLa and SH-SY5Y cells.

### Intramuscular virus injection of neurotropic EV-D68 causes paralysis in neonatal mice

US/MO/47, US/TN, and VR1197 were assessed *in vivo* for their ability to cause paralysis and neuropathogenesis in two-day-old outbred Swiss Webster mouse pups as previously described in Hixon *et. al* [21]. Intramuscular injection of US/MO/47 resulted in limb paresis and paralysis in all mice injected (n=10) as quantified by a Motor Impairment Score (**Fig 2A**) Materials. Most mice injected with US/MO/47 developed moderate to severe paralysis in both rear limbs. Paralysis always began in the injected hind limb and then spread to the contralateral hind limb in most animals, with rare spread of paralysis to the fore limbs. Quantification of the average motor impairment over time showed onset of weakness starting at approximately 4 days post-infection (dpi) with progressive worsening through 7 dpi, with the majority of mice continuing to have moderate to severe weakness in both hind limbs through the end of the observation period at 14 dpi. These data are consistent with previously published results on EV-D68-induced paralysis [21, 43]. In contrast to US/MO/47, mice receiving intramuscular injection of US/TN (n=11) or VR1197 (n=10) failed to develop any signs of motor impairment during the two-week observation period.

**Fig 2.**
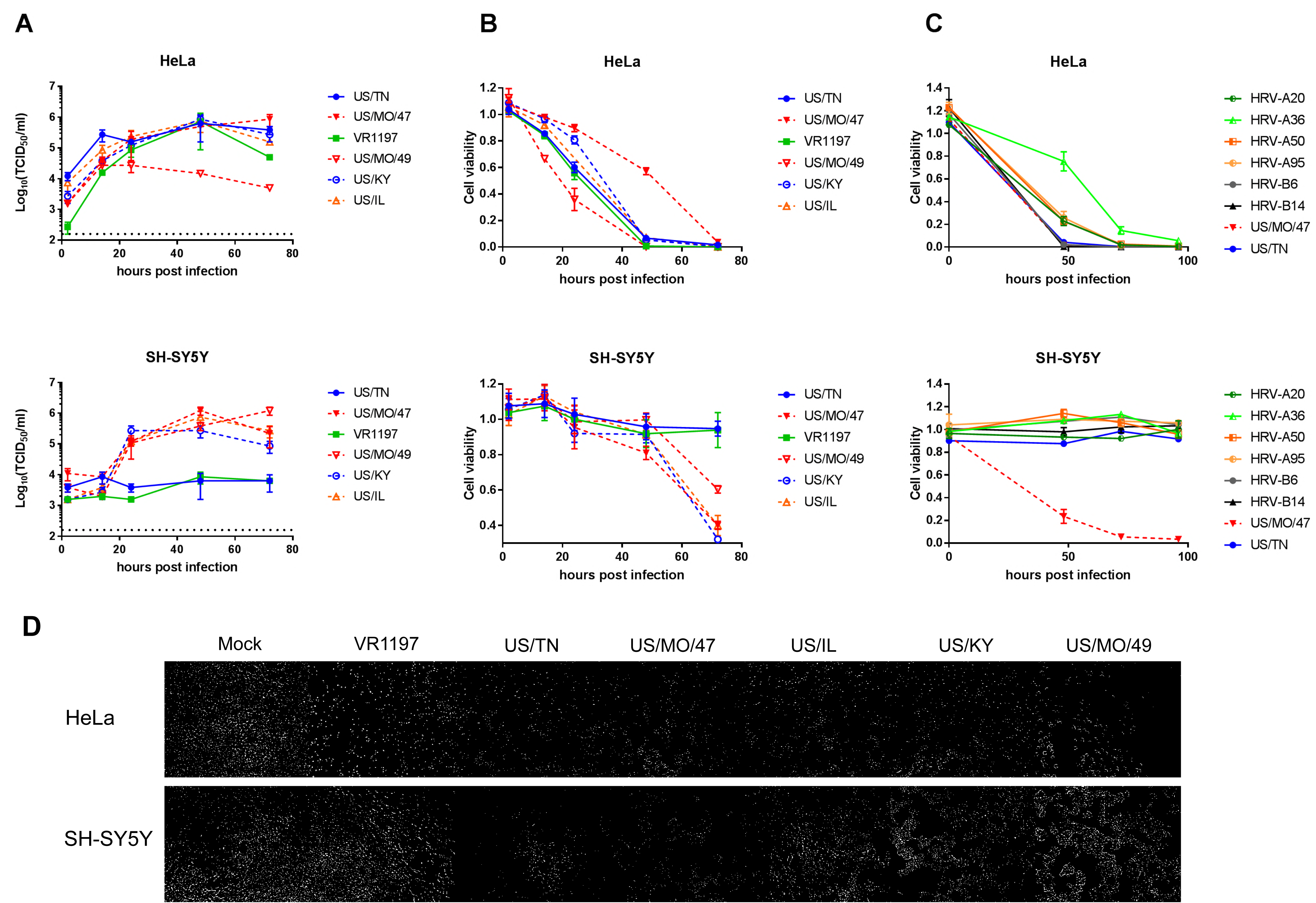
Differential motor impairment in mice following intramuscular injection. A) Motor impairment was scored daily for 14 days post-intramuscular challenges with the indicated EV-D68 strains. None of the mice infected with US/TN or VR1197 mice developed signs of paralysis, whereas 100% of mice infected with US/MO/47 developed paralysis. Error bars represent standard error of the mean. B) Viral titers from muscle and spinal cord titers were determined by TCID_50_ assay on samples taken at 0, 2, 4, and 6 days post-intramuscular infection. Error bars represent the standard deviation.

### Neurotropic EV-D68 can be detected in the spinal cords of paralyzed mice

Intramuscular infection with US/MO/47 resulted in increased titers of infectious virus within mouse spinal cords paralleling the onset of motor impairment. Viral replication was first detected at 2 dpi in spinal cords (∼10^3^ TCID_50_/spinal cord), which corresponded with a rapid increase in viral titer within the muscle tissue (∼10^5^ TCID_50_/mg) of the injected limb (Fig **2B**). Viral titer remained detectable in both spinal cords (∼10^3^ TCID_50_/spinal cord) and muscle tissue (∼10^4^ TCID_50_/mg) at 6 dpi in US/MO/47-injected mice. In contrast, neither US/TN or VR1197 produced detectable infection within mouse spinal cords. We observed sustained viral titers from US/TN within mouse muscle up to 6 dpi (∼10^3^TCID_50_/mg), but no detectable spread of virus to spinal cord. VR1197 did not produce a sustained infection in mouse muscle, and viral titer dropped to the limits of detection by 6 dpi.

### Replication kinetics of recently circulating EV-D68 strain in SH-SY5Y cell culture model

To further characterize the differential replication observed for diverse contemporary EV-D68 isolates, and to test the SH-SY5Y cell infection mode, we obtained all additional commercially available strains of EV-D68. These included strains from the B1, B2, and D1 clades. Interestingly, all additional viral strains replicated in both HeLa and SH-SY5Y (**3A Fig**), including another strain from the B1 clade, US/MO/49, a strain from the newly defined D1 clade (US/KY), and a strain from the B2 clade (US/IL), replicating to a viral titer of ∼10^5^ TCID_50_/ml by 48 hpi. These three strains showed CPE after infection of SH-SY5Y cells (**3D Fig)**. All EV-D68 strains replicated at similar rates in HeLa cells.

To validate our qualitative CPE evaluation, we performed an independent assay of cell death using ATP content as determined by CellTiter Glo luminescence assay (Promega) as a surrogate for viable, intact cells. Using a MOI of 0.1, cell viability dropped after 12 hpi when HeLa cells were infected with every EV-D68 strain tested and continued to drop until the limit of detection was reached, between 48 and 72 hpi (**Fig 3B**). In contrast, cell viability of infected SH-SY5Y cells, beginning at approximately 48 hpi, only dropped for strains where CPE was present. The results of the cell viability and CPE assays was reflective of the TCID_50_ data in HeLa and SH-SY5Y cells for the new panel. The cell viability assay was also performed at 37°C, which produced a similar replication pattern despite initials reports that EV-D68 grows poorly at 37°C [44]. We observed similar rates of viral replication in HeLa and SH-SY5Y cells at both 33°C and 37°C (**S3 Fig**).

**Fig 3.**
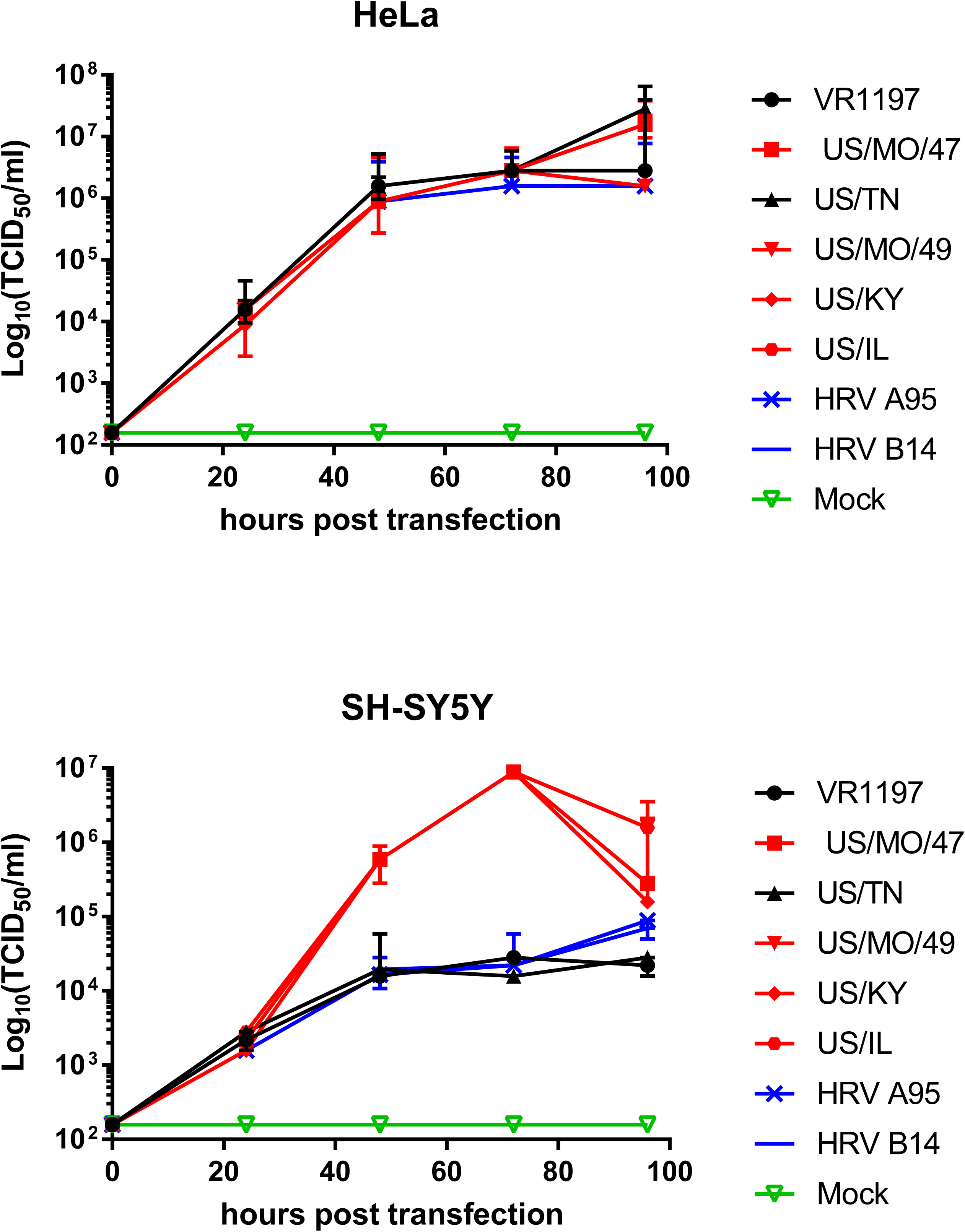
An expanded set of contemporary EV-D68 strains infect SH-SY5Y, but HRV strains do not. A) Hela and SH-SY5Y cells were infected with 6 different EV-D68 isolates at a MOI of 0.1. Cell culture lysates were collected at various time points and viral titer determined by TCID_50_ in HeLa cells. Dotted black lines indicate the limit of detection. Error bars represent SEM of three biological replicates. B) Similarly, cell viability was determined by quantifying the ATP content of the supernatant with CellTiter Glo (Promega) luminescence. Cell viability was calculated relative to mock-infected cultures. Error bars represent SEM of three replicates. C) Six different human rhinovirus (HRV) strains and two EV-D68 strains were used to infect HeLa and SH-SY5Y cell cultures at a MOI of 0.1 and were visualized at 72 hpi. Cell viability was calculated as above. D) Differential cytopathic effects of different EV-D68 isolates in HeLa and SH-SY5Y cells after infection with different EV-D68 isolates at a MOI of 0.1 before visualization at 72 hpi.

### Human Rhinovirus does not infect SH-SY5Y

SH-SY5Y cells were tested for infectivity across a broad selection of HRV strains: two strains from the HRV-B lineage (HRV-B6 and HRV-B14) and four strains from the HRV-A lineage (HRV-A95, HRV-A50, HRV-A36 and HRV-A20). Using the cell viability assay with a MOI of 0.1, the number of viable cells dropped for all HRV strains in HeLa cells. However, in SH-SY5Y cells, no evidence of HRV infectivity was observed for any strain tested using either the cell viability assay (**Fig 3C**) or visual inspection for CPE (**S4 Fig**).

### Differential infection by EV-D68 viral strains is the same in differentiated and undifferentiated SH-SY5Y cells and in primary human neurons

To further characterize the differential replication capability of different EV-D68 strains, we differentiated SH-SY5Y using a well-established retinoic acid (RA) treatment protocol [45, 46], before virus infection, and confirmed differentiation by microscopic examination for morphological changes. We observed no difference in EV-D68 infectivity pattern between differentiated and undifferentiated SH-SY5Y cells. CPE observation and viral replication rate (**S5 Fig**) were similar compared to undifferentiated SH-SY5Y for all strains. All strains that could replicate in undifferentiated SH-SY5Y could also replicate in differentiated SH-SY5Y cells, and viral strains that could not replicate also did not replicate in differentiated SH-SY5Y cells. This demonstrates that EV-D68 strains are capable of infecting neuronal precursors and can also infect mature differentiated neuronal cells. Primary human fetal brain-derived neurons were cultured and infected with US/TN, VR1197 and US/MO/47 (**S6 Fig**). Neurotropic and non-neurotropic EV-D68 strains showed the same infectivity pattern in SH-SY5Y. Using a MOI of 0.01, US/TN and VR1197 plated onto primary neuronal cells did not replicate and viral titers did not rise above the inoculation level baseline. In contrast, an increase in viral titers was observed when the neurotropic strain US/MO/47 infected primary neuronal cells and reached peak titer ∼10^5^ TCID_50_/ml at 24 hpi.

### All EV-D68 strains generate virus when transfected into SH-SY5Y cells

To determine if virus cell entry may be responsible for restricting virus replication in SH-SY5Y for some isolates, full-length genomic RNA from each of the EV-D68 isolates was transfected into the cytoplasm of SH-SY5Y cells and virus production was measured. RNA transfection into HeLa cells resulted in a viral infection and replication pattern similar to intact virus infections at a MOI of 0.1, with viral titers peaking at ∼10^7^ TCID_50_/ml for all EV-D68 strains tested. In contrast to what we observed using standard infection assays, all tested D68 strains generated virus following RNA transfection into SH-SY5Y cells. However, the viral titer peaked at 10^4^ TCID_50_/ml 48 hours after transfection of SH-SY5Y for viral strains that could not infect SH-SY5Y cells using intact virions (**Fig 4**), whereas the viral titer continued to increase until saturation at approximately 10^7^ TCID_50_/ml following transfection of SH-SY5Y with strains that could infect SH-SY5Y cells.

**Fig 4.**
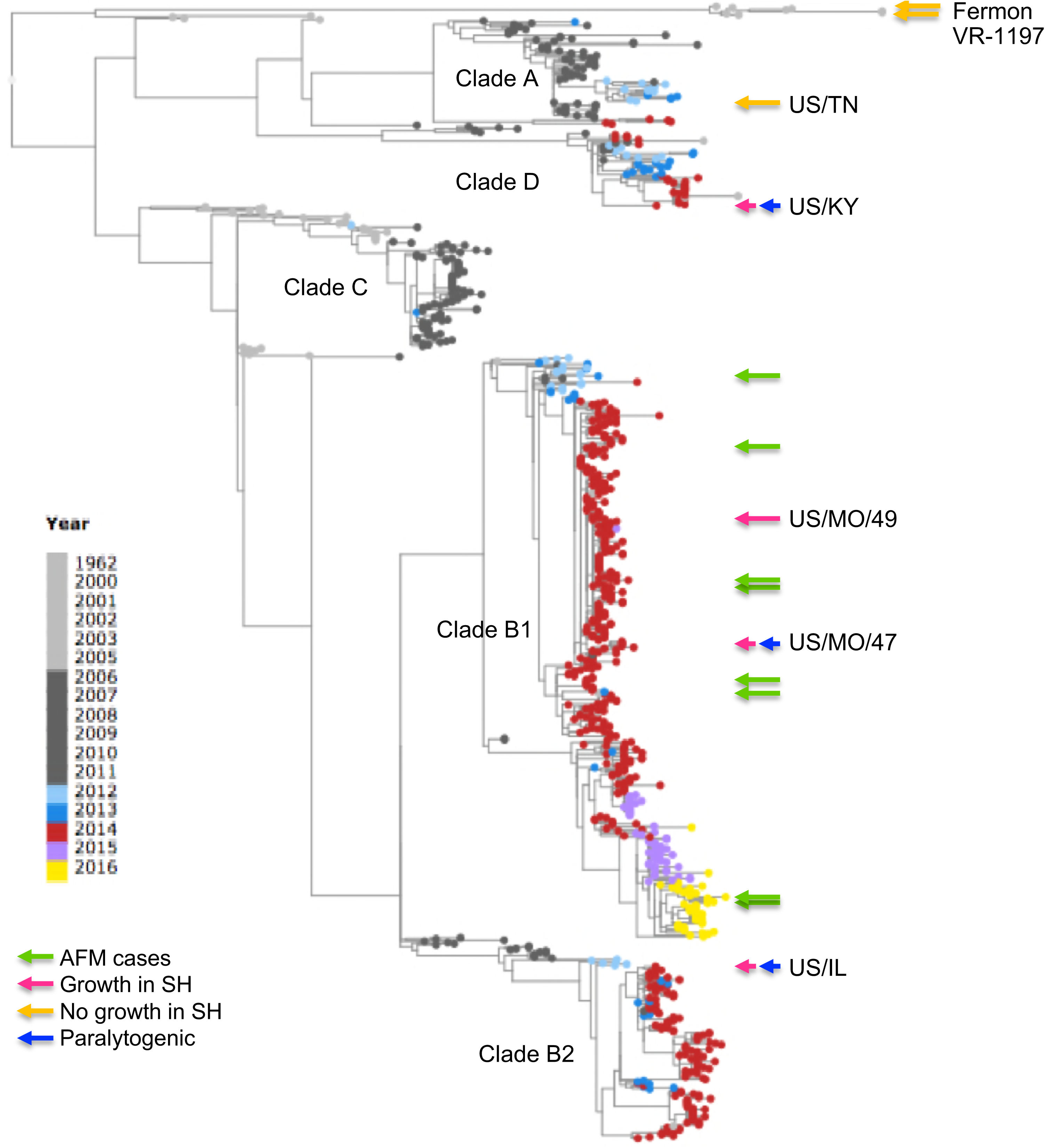
Replication of EV-D68 isolates following transfection of genomic RNA into HeLa and SH-SY5Y cells. RNAs were purified from various HRV and EV-D68 virus stocks and used to transfect SH-SY5Y and HeLa cells. Cell culture lysates were collected at various time points and viral titer determined by TCID_50_ in HeLa cells. Error bars represent SEM of three biological replicates.

## Discussion

Here we report on the differential infectivity between various contemporary and historical EV-D68 strains in SH-SY5Y as measured by viral replication, cell viability, CPE, and immunofluorescence. The clade specificity of neuropathogenesis previously reported [10, 22, 25-27, 32, 33] between contemporary and historical strains is observed in the neurotropism in SH-SY5Y cells (**Figure 5**). Among the EV-D68 strains used in this study, those from clades B1, B2 and D1 were able to infect SH-SY5Y cells and cause paralysis in a mouse model, whereas those from clade A and other historical strains could not [47]. We also showed that this differential growth is at least partly due to differential viral entry, as all strains can replicate and produce viral progeny after transfection.

**Fig 5.**
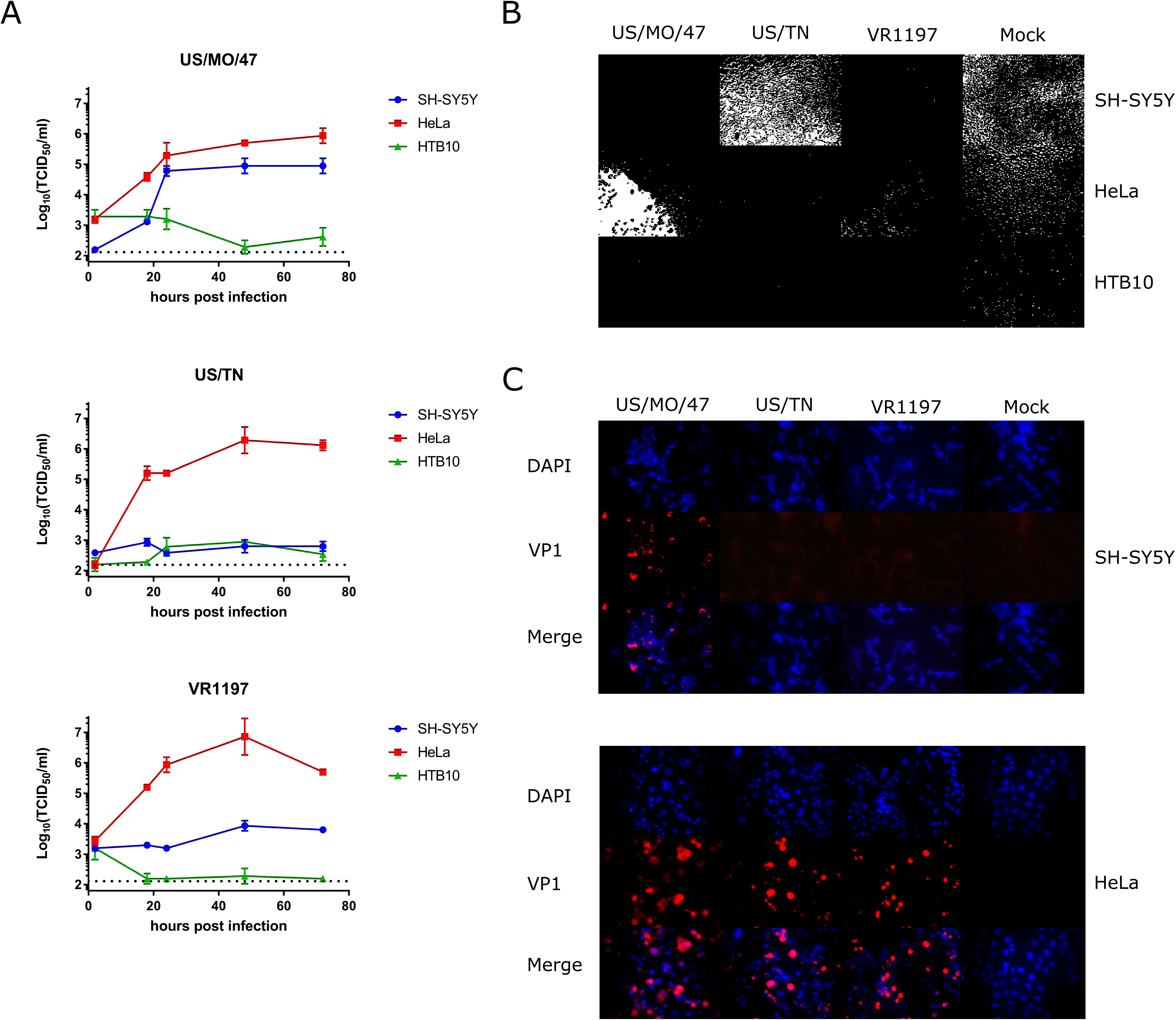
Phylogenetic tree of EV-D68 isolates based on VP1 sequences. VP1 nucleotide sequences of > 900 nt were retrieved from the ViPR site (https://www.viprbrc.org/brc/home.spg?decorator=picorna_entero)” www.viprbrc.org/brc/home.spg?decorator=picorna_entero) on July 24, 2017. Sequences were aligned using the MUSCLE algorithm and sequences showing poor alignments were removed. A phylogenetic tree was computed using RaXML (bootstrap replicates of 100) and then visualized using Archaeopteryx.js, via the ViPR site. Clade classifications are based on bootstrap values of 100%. AFM-associated isolates are marked with a green arrow. EV-D68 isolates used in this study are labeled either with a pink (growth in SH-SY5Y cells) or orange (no growth in SH-SY5Y cells) arrow. Isolates that are paralytogenic in mice (56) are labeled with a blue arrow.

To determine the most appropriate tissue culture model system, we explored the gene expression by RNA sequencing and showed that transcripts from SH-SY5Y cells are enriched for neuronal-specific genes relative to HeLa and HTB10 cells. This agrees with several other groups which have shown the expression of specific neurological marker genes in SH-SY5Y cells [45, 48, 49]. This neuronal cell line has successfully been used as a model to study the *in vitro* neuropathogenic effects of different viruses, notably, as a model for paralytic enteroviral isolates including EV-A71 and poliovirus [35, 50-56]. Numerous other viruses associated with neurological symptoms can infect SH-SY5Y, including: Japanese encephalitis-virus [57], human immunodeficiency-virus [58], human cytomegalovirus [59], varicella-zoster virus [60], chikungunya [61], mumps virus [62], dengue virus [63], Zika virus [64, 65], and rabies virus [66] suggesting that SH-SY5Y is an excellent model system for neurotropic viruses.

We found support of the biological relevance of SH-SY5Y cells as a cell culture model by observing the same infectivity pattern *in vitro* with other neuronal cell cultures. SH-SY5Y cells are capable of being differentiated using retinoic acid, which leads to more neuron-specific morphology and gene expression. These differentiated cells are characterized by the formation of extensive neurites, as well as the induction of neuron-specific enzymes, receptors and neurotransmitters [45]. Differentiated SH-SY5Y cells have been used as models for neuron-virus interactions [67] including EV-A71 [55, 56], Chikungunya virus [61], and varicella-zoster virus [60]. Our results show that both differentiated and undifferentiated SH-SY5Y can be infected with neurotropic strains of EV-D68, suggesting neurotropic strains of EV-D68 can invade both immature and mature neuronal cells. Primary human fetal brain-derived neurons have also been used to study infectivity patterns of neurotropic viruses as well as the central nervous system [68, 69]. We also observed a similar EV-D68 replication pattern in human primary postnatal neurons. Interestingly, despite initially reported as growing poorly at 37°C [44], we observed similar rates of EV-D68 replication in SH-SY5Y cells at both 33°C and 37°C, which would likely be a more biologically relevant temperature for paralytic infections at the core body temperature rather than the lower temperature of the upper respiratory system that supports more routine respiratory infection.

EV-D68 is closely related to human rhinovirus (HRV) within the *Picornaviridae* family [70]. Frequent coinfection in patients and cross-reactivity of nucleic acid amplification screening had led to misdiagnosis of EV-D68 infection as HRV infection prior to the 2014 outbreak [71, 72]. Despite the similarity in respiratory symptoms, HRV lacks any association with the neurological symptoms of other enteroviruses, such as EV-D68 and EV-A71 [70]. Indeed, despite their similarity to EV-D68, none of the 6 HRV strains could infect SH-SY5Y cells.

We hypothesized that the reason for differential replication in SH-SY5Y cells of EV-D68 strain relates to different viral entry capabilities. We used RNA transfection to deliver infectious RNA to the cytoplasm, bypassing natural viral entry mechanisms during an infection. All EV-D68 strains generated virus following RNA transfection, but in some cases viral titers plateaued at a relatively lower level, suggesting that only a single round of replication had occurred. HRV RNA transfected into SH-SY5Y cells produced a similar result. Our interpretation is that differences in the sequence and structure of viral capsid proteins are responsible for the differential infectivity in SH-SY5Y cells by EV-D68 strains, and that viral entry is what prevents HRV, US/TN, and the historical strains from infecting SH-SY5Y cells. It has recently been reported that a chimeric swap mutant exchanging the viral capsid from EV-D68 VR1197 and a neurotropic EV-D94 strain capable of replication in SH-SY5Y cells, results in a loss on infectivity in SH-SY5Y cells [47]. This result further supports our conclusion that viral entry mediated by the capsid is the cause of the observed differential neurotropism. Specific genetic residues may be the cause of differential neurotropism. A comparative analysis using infectious clones bearing specific polymorphisms will likely be needed to establish the determinants of neurotropism in SH-SY5Y. In particular, the 2014 outbreak B1 substitutions in VP1/98A, VP1/148V, VP1/280K, VP1/290S, VP1/308N, and VP2/222T are all located on the virion surface and could be directly involved in virus-host cell attachment and would be good candidates to evaluate.

Our results closely correlate with the differential paralytic myelitis caused by EV-D68 in mice suggesting that infectivity in SH-SY5Y cells may be an effective proxy for neuropathogenesis (**Figure 5**). Multiple contemporary EV-D68 strains that have been shown to cause paralytic myelitis in mice were also neurotropic in SH-SY5Y cells [21]. The historical, non-neurotropic strains that are nonparalytic in mice did not grow in SH-SY5Y cells. In particular, the contemporary EV-D68 strain, US/TN, which was non-neurotropic in SH-SY5Y cells and not previously reported in *Hixon et al*. failed to produce paralysis in mice and could not be found in mouse spinal cord tissue. US/TN appeared to replicate at a low level within mouse muscle tissue, indicating lack of paralysis was not due to an inability to infect the mice. The infectivity pattern in SH-SY5Y cells, along with the agreement in primary postnatal neurons, supports the mouse model reported in Hixon *et al.* [21] and validates the results in the SH-SY5Y human cell line. This is significant because, while useful, mouse models are costly and introduce potential caveats, such as transcriptional factor differences in mice versus humans [73-75] and is debated if the mouse model recapitulates human conditions [76, 77]. Evidence presented here supports the validity of the mouse model for further study of neurotropic EV-D68 viruses and the theory of a causal link between AFM in humans and EV-D68.

## Conclusion

We present a differential neuronal infectivity phenotype between contemporary and historical EV-D68 strains. Permissible infection of SH-SY5Y cells mimics the paralysis pattern reported in animal models. The high throughput nature of tissue culture models will allow for rapid screening of novel viral strains and recombinant viruses to elucidate the genetic determinants of neurotropism and potential antiviral therapies. This can enable identification of EV-D68 alleles responsible for neural infection and potentially neurological disease and avoids the cost associated with large-scale screening using animal models.

## Materials and Methods

### Ethics Statement

All studies were done in accordance with the University of Colorado IACUC and Animal Use Committee (B-34716(03)1E). Mice were cared for in adherence to the National Institute of Health (NIH) guidelines to the Care and Use of Laboratory Mice. Mouse pups exhibiting paralysis were euthanized if unable to nurse. Mice were anaesthetized with inhaled isoflurane before tissue collection or perfusion.

### Cell culture

HeLa cells (ATCC) were maintained in Dulbecco’s modified Eagle’s medium (DMEM, Gibco) supplemented with 10% fetal bovine serum (FBS, HyClone). HTB10 (ATCC) cells were maintained in DMEM supplemented with 10% FBS and non-essential amino acids (Gibco). SH-SY5Y (CLR-2266, ATCC) cells were maintained in a 1:1 mixture of DMEM and F12 (Gibco) media supplemented with 10% FBS. To differentiate SH-SY5Y cells, a ∼50% confluent flask of SH-SY5Y cells had media replaced with 1:1 mixture of DMEM and F12 (Gibco) media supplemented with 3% FBS and 10 µM retinoic acid (RA, Sigma) [53]. After 3 d of exposure to RA, the morphology of cells was evaluated, and cells were passaged for further use. Morphology and cytopathic effect was evaluated using an inverted microscope. Human postnatal day 0 (P0) brain neurons were purchased from ScienCell (Cat# 1520) and plated at a density of 40,000 neurons/well on poly-D-lysine coated 96-well plates [69]. The neurons were maintained in ScienCell neuronal growth media with penicillin-streptomycin (Cat# 1520) at 37°C in 5% CO_2_ until day in vitro (DIV) 7, by which time the neurons had well-established neurites.

### Virus Stock Preparation

EV-D68 stocks were prepared by infecting HeLa or rhabdomyosarcoma cells (ATCC) at 33°C in 5% CO_2_ until CPE was observed. Cell debris was removed by centrifugation and titers determined in a standard Tissue Culture Infective Dose (TCID_50_) assay and calculated by the Spearman-Kärber method. The source of each strain is detailed in (**S1 table**)

**Table S1:**
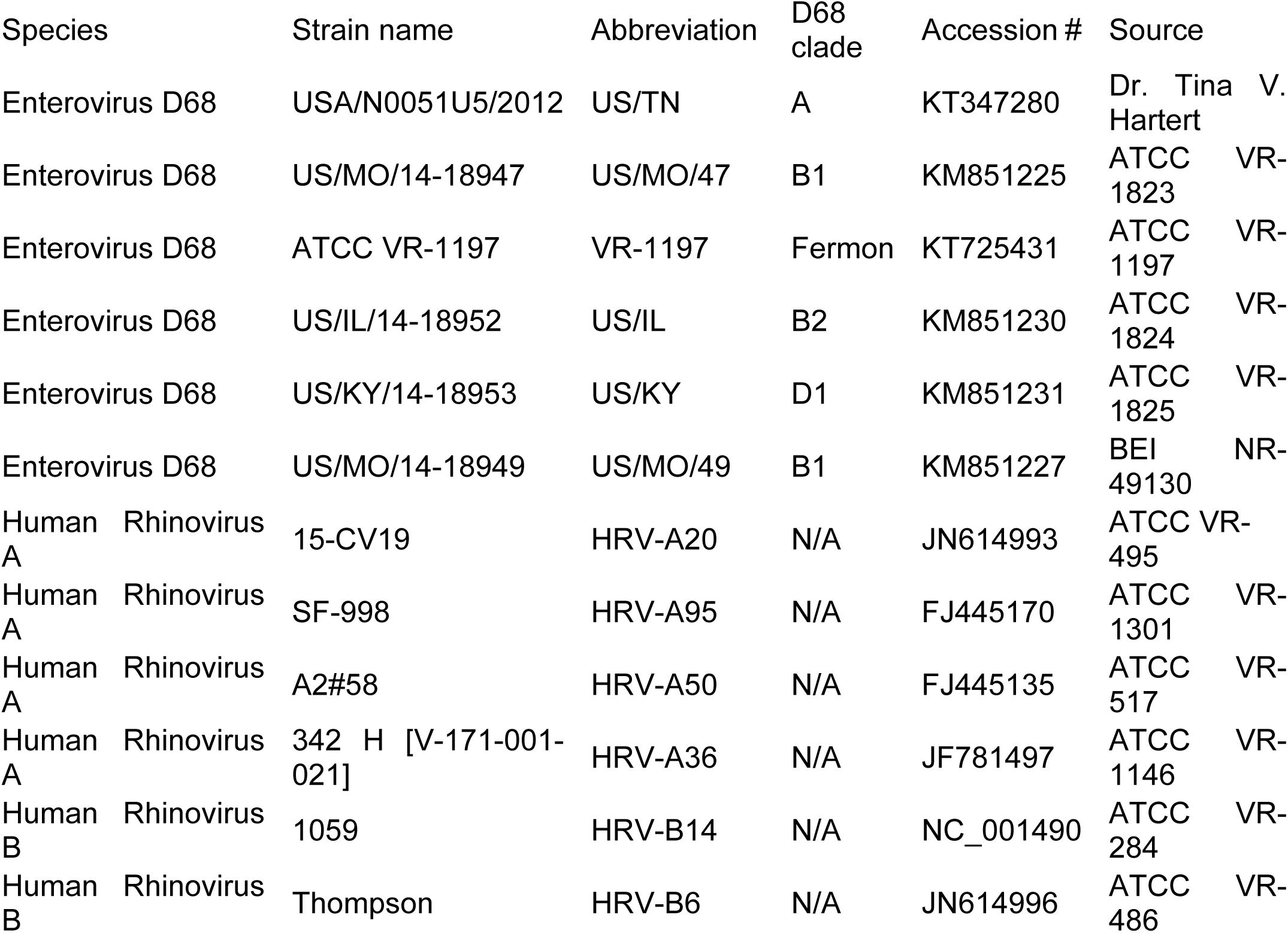
List of strains used in this study.

| Species | Strain name | Abbreviation | D68<br>clade | Accession # | Source |
| --- | --- | --- | --- | --- | --- |
| Enterovirus D68 | USA/N0051U5/2012 | US/TN | A | KT347280 | Dr. Tina V. Hartert |
| Enterovirus D68 | US/MO/14-18947 | US/MO/47 | B1 | KM851225 | ATCC VR-1823 |
| Enterovirus D68 | ATCC VR-1197 | VR-1197 | Fermon | KT725431 | ATCC VR-1197 |
| Enterovirus D68 | US/IL/14-18952 | US/IL | B2 | KM851230 | ATCC VR-1824 |
| Enterovirus D68 | US/KY/14-18953 | US/KY | D1 | KM851231 | ATCC VR-1825 |
| Enterovirus D68 | US/MO/14-18949 | US/MO/49 | B1 | KM851227 | BEI NR-49130 |
| Human Rhinovirus A | 15-CV19 | HRV-A20 | N/A | JN614993 | ATCC VR-495 |
| Human Rhinovirus A | SF-998 | HRV-A95 | N/A | FJ445170 | ATCC VR-1301 |
| Human Rhinovirus A | A2#58 | HRV-A50 | N/A | FJ445135 | ATCC VR-517 |
| Human Rhinovirus A | 342 H [V-171-001-021] | HRV-A36 | N/A | JF781497 | ATCC VR-1146 |
| Human Rhinovirus B | 1059 | HRV-B14 | N/A | NC_001490 | ATCC VR-284 |
| Human Rhinovirus B | Thompson | HRV-B6 | N/A | JN614996 | ATCC VR-486 |

### Replication kinetics of EV-D68

The replication kinetics for HeLa, A549, HTB10, SH-SY5Y and differentiated SH-SY5Y cell were evaluated in a high throughput manner. Viral replication kinetics were measured from sets of flat bottom 96-well plates. Sets of plates corresponding to the number of desired timepoints in an experiment were infected at the same initial time, using distinct 96-well plates for each time point. Infected plates were incubated at 33-34°C, 5% CO_2_ until the designated time point when each corresponding plate was placed in a −80°C freezer until the entire time course was completed. Mock infected wells adjacent to each condition demonstrated that no contamination occurred across wells. After 2 h, high MOI conditions (0.1 and 1) were washed three times with phosphate buffered saline (PBS) and the 2 h time point plate frozen to determine the background levels of virus present, since 2 h is long enough for EV-D68 entry but not long enough for replication. After three freeze-thaw cycles the viral titers from 10-fold serial dilutions of each sample were evaluated using a 50% TCID_50_ assay on HeLa cells. Plates were scored after adding 100µl of crystal violet fixative per well followed by a 1 h incubation at room temperature (RT) and washing to remove unbound dye. Crystal violet fixative was prepared by adding 5 g crystal violet (sigma) and 8.5 g sodium chloride (Sigma) to 50 ml formaldehyde, 260 ml ethanol, and 690 ml deionized water.

For replication kinetics in human postnatal neurons, day in-vitro (DIV) 7 neurons were infected with EV-D68 US/MO/47, US/TN, or VR1197 at a MOI = 0.01. Infection media was left on the cells for the duration of the experiment to minimize loss of cells from multiple rinses due to low cell adhesion. Cell culture supernatant and lysate was collected at 0, 6, 12, 24, 48, and 72 h, with three biological replicates collected per time point for each viral strain. Lysate was serially-diluted 10-fold from 1 (raw lysate) to 10^−6^ RD cells at 33°C and evaluated using a 50% TCID_50_ assay.

### Immunostaining

HeLa and SH-SY5Y cells were grown to 50-70% confluence on coverslips in a 24-well plate and infected with EV-D68 strains at a MOI of 1.0. Mock infected cells serve as a negative control. Coverslips were incubated at 34°C, 5% CO_2_ for 18 hours, then fixed with 4% paraformaldehyde (PFA) and stored at 4°C. The coverslips were washed with PBS and cells were permeabilized with 0.1% Triton-X for 10 minutes. The coverslips were blocked with 2% bovine serum albumin in PBS for 1 hour. The cells were incubated with rabbit polyclonal α-VP1 of EV-D68 (GeneTex) at a final concentration of 4 µg/ml overnight at 4°C, washed 3 times, and then incubated with a secondary goat α-rabbit rhodamine red-X (Thermo Fisher) at a final concentration of 1 µg/ml for 30 minutes. To visualize nuclei, DAPI stain was added to the second of three washes. The wells were visualized on an Axioskop 2 plus (Zeiss) fluorescence microscope using DAPI and Rhodamine filters. Images were taken with AxioCam MRc5 (Zeiss) camera using AxioVision software. All images for a particular filter were taken under identical exposure conditions.

### Mouse Infections with EV-D68

Animal experiment were performed in an AAALAC accredited animal facility under IACUC protocol B-34716(03)1E at the University of Colorado. Pregnant female Swiss Webster mice were ordered from Envigo and kept in standard housing until the pups were born. At post-natal day 2, the dam and pups were transferred the BioSafety Level 2 (BSL2) region of the animal facility. P2 Swiss Webster mouse pups were then inoculated with 10^6^.^8^ TCID50/ml virus in 10 ul by intramuscular injection into the left medial hindlimb [21].

Mouse pups of both sexes were randomized to treatment conditions before virus inoculation.

### Motor Impairment Scoring

Mice were monitored daily for 14 days. To assess paralysis, mice were removed from the cage and observed moving on a flat surface for several minutes in which each limb was given a motor impairment score: 0 - no motor impairment; 1 - mild motor impairment, ataxia or decreased movement present, toe/knuckle walking; 2 - moderate impairment, profound ataxia, limited movement of limb; 3 - severe impairment, no movement in limb, limb is non-weight bearing. The final motor impairment score for each day was achieved by summing the score for each limb.

### Mouse tissue collection

Mouse pups were sacrificed by decapitation for collection of muscle and spinal cord tissue. Spinal cords were removed as previously described [21, 43]. Muscle tissue was collected from the inoculated limb (with the goal of obtaining as much muscle tissue possible from the anterior and posterior thigh and gastrocnemius). Both tissues were collected into BeadBug tubes containing inert ceramic bead and 0.3 mL of ice-cold, sterile PBS. Tissues were lysed mechanically on a BeadBug tissue homogenizer for 45 seconds at 2800 rpm, and stored at −80°C. After thaw, tissue samples were spun at 2700xg for 1 minute to remove tissue chunks from the lysate. Lysate was serially-diluted 10-fold from 1 (raw lysate) to 10^−6^ and plated in a standard TCID_50_ assay to determine the final viral titer. To get the final titer per whole spinal cord, TCID_50_/mL was multiplied by 0.3 mL. To get the final muscle titer per milligram of tissue, TCID_50_/mL was multiplied by 0.3 mL and divided by the weight of tissue collected. Samples that were below the limit of detection were graphed at zero.

### Cell ATP/viability assay

Cells were cultured and evaluated in the same manner as in the viral replication kinetics assays. ATP levels were measured using the CellTiter-Glo luminescent cell viability assay kit (catalog number G7570; Promega), and cell viability calculated relative to mock control. To preserve the ATP levels so that each timepoint could be evaluated concurrently, cell supernatant was removed and cells frozen at −80°C. Once the time series was completed, all plates were removed and RT media was added to each plate. Upon stabilization at RT for 20 min, the manufactures protocol was followed. We validated this deviation from the manufactures protocol by confirming the linearity of the assay across the active range of the study.

### RNA sequencing

In order to explore the use of SH-SY5Y as an appropriate neuronal cell model, we used RNA sequencing to obtain a comprehensive view of the genes expressed in SH-SY5Y, HeLa and HTB10 cell lines. To prepare cells for RNA sequencing 10^4^ cells were grown in a 96 well plate for 24 h in quadruplicate before washing and resuspension in 10 µl of Cell Lysis Buffer (0.2% Triton X-100, 2 Units/µL RNase inhibitor, 1:2,000,000 dilution of ERCC spike-in RNAs (Life Technologies)) per well. Full length cDNA was amplified using the SmartSeq2 protocol optimized in our laboratory [78, 79] before Nextera XT library preparation and sequencing on a NextSeq500 with 2 × 150 paired end reads. After adapter/primer trimming using the Trimmomatic tool (http://www.usadellab.org/cms/?page=trimmomatic), trimmed sequencing reads were mapped to transcripts derived from the human reference genome (GRCh37) and gene expression levels (transcripts per million reads) estimated using the RSEM package [80].

### RNA purification and transfection

RNA used for transfections was purified from viral stocks grown in HeLa cells. Purification was performed using QIAamp MinElute Virus Spin Kit (Qiagen) according to the manufacturer’s instructions; final RNA concentrations were approximately 100 ng/µl. In 12-well plates selected cell cultures were seeded and grown. According to the manufacturer’s instructions, 200 ng of RNA was used with 2 µl of each reagent in the *Trans*IT®-mRNA transfection kit (Mirus) to perform a transfection.

## Acknowledgements

We thank Dr. Tina V. Hartert and her colleagues at Vanderbilt for US/TN EV-D68 strain.

## Supporting Information

**S1 Table. List of strains used in this study.**

**S1 Fig. EV-D68 virus titers in A549 cells.** A549 cells were grown to 90% confluence in 96 well plates before infection with EV-D68 US/MO/47, US/TN and VR1197 EV-D68 at a MOI = 0.1. Infection media was removed after 2 hpi to reduce background. Cell culture lysate/supernatants were collected at various time points after infection, and viral titers measured using endpoint dilutions for growth in HeLa cells. Dotted black line indicates the limit of detection. Error bars represent SEM of three biological replicates.

**S2 Fig. EV-D68 virus titers in three different cell cultures with additional MOIs.** Cells from three different cell lines – SH-SY5Y, HTB10, and HeLa were grown to 90% confluence in 96 well plates before infection with EV-D68 US/MO/47, US/TN and VR1197 EV-D68 at a MOI = 1.0 and MOI = 0.01. Infection media was removed after 2 hpi to reduce background from MOI = 1.0. Cell culture lysate/supernatants were collected at various time points after infection, and viral titers measured using endpoint dilutions for growth in HeLa cells. Dotted black line indicates the limit of detection. Error bars represent SEM of three biological replicates.

**S3 Fig. Cell viability in cells infected with EV-D68 at 37°C.** Using replicate plates, cell viability was measured by quantifying ATP content as determined by CellTiter Glo (Promega) luminescence. Cell viability calculated relative to mock. Error bars represent SEM of four replicates.

**S4 Fig. HRV does not infect SH-SY5Y.** Six different HRV strains and two EV-D68 strains were used to infect HeLa and SH-SY5Y cell cultures grown in a 96-well plate at a MOI of 0.1 before visualization at 72 hpi.

**S5 Fig. EV-D68 virus titers in differentiated SH-SY5Y cells.** Differentiated SH-SY5Y were infected with 6 different isolates of EV-D68 at a MOI of 0.1. Cell culture lysate/supernatants was collected at various time points. Viral titer was determined by TCID_50_ in HeLa cells. Dotted black line indicates the limit of detection. Error bars represent SEM of three biological replicates. Error bars represent SEM of three replicates.

**S6 Fig. EV-D68 strain growth in human postnatal cortical neurons.** Human postnatal day 0 brain neurons were maintained to day 7 *in vitro* before infection with EV-D68 US/MO/47, US/TN, or VR1197 at a MOI = 0.01. Cell culture lysates/supernatant were collected at various times post viral infection, and viral titers were measured using endpoint dilutions for growth in RD cells. The x-axis indicates the limit of detection. Error bars represent SD of three biological replicates.

